# Host-independent persistence of the *Diadema antillarum* Scuticociliatosis *Philaster* clade in coastal environments

**DOI:** 10.64898/2026.01.07.698123

**Authors:** Michael W. Henson, Axel J. Leon-Rodriguez, Christopher M. DeRito, Harilaos Lessios, Jean-Pascal Quod, Brayan Vilanova-Cuevas, Ian Hewson

**Affiliations:** Department of Biological Sciences, Northern Illinois University, DeKalb IL USA; Department of Microbiology, Cornell University, Ithaca NY USA; Smithsonian Tropical Research Institution, Balboa, Panama; ARVAM, Réunion, France

**Author notes:** Correspondence: Michael W. Henson, Ian Hewson.

## Abstract

The mass mortality of the keystone herbivore *Diadema antillarum* in the Caribbean was caused by the pathogenic ciliate from the *Diadema antillarum* Scuticociliatosis *Philaster* clade (DaScPc). Despite its confirmed pathogenicity, the environmental distribution and persistence strategies of DaScPc outside its host remain poorly understood. We used quantitative PCR, nested PCR, and Sanger sequencing across a 16-month time series and broad geographic surveys to investigate its ecological dynamics and potential environmental reservoirs. Sequencing-confirmed detections at a Florida coastal site devoid of *D. antillarum* show that DaScPc is a natural, host-independent component of the reef environment. Molecular detection on coral and macroalgal surfaces and in the plankton fraction indicates that multiple substrates can harbor the ciliate. Temporal observations revealed emerging trends with macroalgal cover and reef productivity, though no direct correlations were observed. Geographically, DaScPc was absent from outbreak sites in Panama and Réunion. Together, this data suggests spatial patchiness and a cryptic “boom-and-bust” lifestyle in which the organism persists at low abundance between outbreaks. The co-occurrence of a related nonpathogenic ciliate (*Acropora/CHN/2009*) further underscores the ecological complexity of the *Philaster* clade. These findings broaden understanding of DaScPc ecology, confirming environmental persistence independent of its host and suggesting that parasitism may be intermittent, triggered by changing environmental conditions. This work highlights the need for higher-resolution surveillance and long-term monitoring to link ecosystem processes with the reemergence of marine disease in vulnerable coral reef systems.

**Importance:** The 2022 mass mortality of the sea urchin *Diadema antillarum* devastated Caribbean coral reefs, yet little is known about how the pathogenic ciliate responsible for the event persists in nature. We show that the *Diadema antillarum* Scuticociliatosis *Philaster* clade (DaScPc) occurs in reef environments even where the host *(D. antillarum*) is absent, indicating that it can persist independently of its host. Our findings suggest that DaScPc occupies a cryptic ecological niche within reef microbial communities and may follow a “boom–bust” dynamic, fluctuating between rare environmental states and occasional proliferation. Although direct environmental drivers remain unresolved, emerging trends with macroalgal cover and reef productivity highlight the potential influence of ecosystem conditions on ciliate abundance. This work broadens the understanding of how marine pathogens persist between outbreaks and underscores the importance of environmental surveillance for predicting and mitigating future disease events on coral reefs.

## Introduction

Marine ciliates are integral components of coastal and pelagic ecosystems, mediating nutrient cycling and energy transfer through the microbial loop via diverse feeding strategies, including bacterivory, herbivory, and detritivory (1–3). Their ecological significance has received increasing attention in recent years, particularly as emergent ciliate taxa have been implicated in mass mortality events among both invertebrates and vertebrates (4, 5). Despite their recognized importance, comparatively little is known regarding the environmental distribution, trophic behavior, and survival strategies of many ciliates outside of their hosts (6, 7). These knowledge gaps are particularly consequential when ciliates act as pathogens, as their persistence and proliferation in the environment may influence both disease emergence and virulence. Detection of ciliates outside their hosts does not necessarily indicate imminent infection, but it raises critical questions regarding the ecological conditions under which pathogenic activity may be triggered.

One pathogenic ciliate of increasing concern is the Diadema antillarum Scuticociliatosis Philaster clade (DaScPc) (8, 9), which caused widespread mortality of sea urchins across the Caribbean in spring and summer 2022 (8, 10). Experimental infection trials fulfilling Koch’s postulates have confirmed its pathogenicity, providing rare direct evidence of a causative agent in a marine disease (8). Field surveys demonstrate that DaScPc is consistently associated with scuticociliatosis in D. antillarum populations throughout the Caribbean (7, 11). The disease appears to be restricted to urchins within the Diadematidae family, with no reports in other co-occurring urchin families (12). Recent studies further indicate that DaScPc may affect a broader range of Diadematidae species across geographically disparate regions, underscoring its expanding ecological relevance (9, 13–20). Beyond its ecological impact, DaScPc represents a rare and valuable system in marine disease research, as the causative pathogen has been unequivocally identified and experimentally validated.

Despite this clarity regarding its pathogenicity, the environmental distribution of DaScPc remains poorly understood. The ciliate has been detected on a range of invertebrates, marine plants, and abiotic surfaces through swab surveys at both currently and recently affected sites (7). Among these detections, corals, particularly the starlet coral (*Siderastrea siderea*), consistently yielded DaScPc sequences, whereas turf algae and the sponge *Ircinia campana* exhibited more sporadic occurrences. These patterns are thought to reflect the availability of dissolved organic matter (DOM) on these surfaces rather than active infection by the ciliate (21). However, the relationships among DOM concentration, bacterial abundance, and DaScPc proliferation have not been empirically investigated. Accordingly, it remains uncertain whether the presence of DaScPc on these substrates represents a potential environmental reservoir or a future risk to D. antillarum or other Diadematidae species (7).

The present study investigates the ecological dynamics of DaScPc through two complementary approaches. First, we conducted temporal sampling at a coastal site near the location where the original DaScPc culture was isolated to test the hypothesis that DaScPc abundance in the plankton and on coral surfaces varies with fluctuations in primary producer biomass and dissolved organic carbon (DOC) concentration. Second, we examined associations between DaScPc and sympatric surfaces, including corals, across two geographically distinct sites: Panama, a region unaffected by the 2022 Diadema antillarum die-off but historically significant as the epicenter of the species’ initial mass mortality event in 1982 (22, 23); and Réunion Island (Western Indian Ocean), the most geographically distant site from the Caribbean where DaScPc-linked scuticociliatosis was reported in autumn 2023 (20). By combining high-resolution temporal sampling with geographically broad surveys, this study aims to improve understanding of DaScPc biogeography, environmental associations, and potential links to mass mortality events.

## Methods

### Assessment of Swab Efficiency for DaScPc Recovery

To evaluate the detection threshold for DaScPc in swab surveys, we conducted an aquarium experiment that mimics natural coral sampling conditions. Twelve fragments of *Astrangia poculata* were maintained in artificial seawater (Instant Ocean) under a 12-hour light/dark cycle. Fragments were transferred individually into 500 mL beakers containing artificial seawater and inoculated in triplicate with 0 (media only), 30, 300, or 3,000 cells of DaScPc culture FWC2.

After a 2-hour incubation, coral surfaces were swabbed following the same procedure used in field collections. Swabs were held in dry transport tubes for 30 minutes to simulate field transport, then fixed in RNAlater for one hour prior to extraction. To compare swab recovery efficiency to known cell counts, we quantified 28S rRNA gene copy number in a dilution series of actively growing DaScPc FWC2 (6–600 cells) using quantitative PCR as described below.

### Temporal Survey of DaScPc and Related Ciliates

A 16-month time series (August 2023–December 2024) was conducted at Sunset Park, Grassy Key, FL, USA (24.759°N, 80.967°W), near the original site where the DaScPc FWC2 culture was isolated (8). The site, characterized by shallow water (0.5–1.2 m) and abundant coral colonies, was sampled every 2–3 months (n = 7).

During each sampling event, 0.6–2.0 L of seawater was filtered through 47 mm, 2.7 µm GF/D filters (Whatman) to capture planktonic ciliates (>2.7 µm). Filters were immediately frozen on dry ice, transported to the laboratory, and stored at –80°C until extraction. Duplicate filtrate samples (50 mL) were collected to quantify inorganic nutrient concentrations (NO₃⁻ + NO₂⁻, NO₂⁻, PO₄³⁻, NH₄⁺, and Si) at the Chesapeake Bay Laboratory Nutrient Analytical Services Laboratory. Water temperature, salinity, pH, and dissolved oxygen were recorded in situ using a handheld YSI ProDSS (Xylem Inc.). Additional filtrate samples were collected into pre-combusted glass vials (1.2 µm GF/F pre-filtration), acidified with 150 µL HCl, and analyzed for dissolved organic carbon (DOC) at the same facility (24).

Concurrent with water sampling, a ∼0.39 km² area was surveyed by two snorkelers for 1–2 hours to collect swabs from coral, sponge, and macroalgal surfaces. Between 18–76 swabs were collected per event (total n = 252; Table 1), depending on local substrate abundance. Each specimen was photographed *in situ* using a GoPro camera for identification. Swabs (Puritan Dry Transport Systems) were rubbed gently over ∼2 cm² of each surface and returned to dry transport tubes. Onshore, swab tips were clipped into cryovials containing RNAlater and transported to Cornell University at ambient temperature for molecular analysis.

**Table 1:** Total number of swab specimens surveyed for the presence of DaScPc by qPCR and conventional PCR.

| Sampling Date | Coral | Porifera |  |  | Macroalgae |  |  |  |  |  |
| --- | --- | --- | --- | --- | --- | --- | --- | --- | --- | --- |
|  | *Siderastrea sideraea* | *Cliona varians* | *Spheciospongia vesparium* | *Tedania ignis* | *Ircinia felix* | *Laurencia* sp. | *Penicillus dunetosis* | *Sargassum* sp. | *Thalassia testudinum* | Other |
| 16-Aug-23 | 23 | 8 | 1 |  |  | 3 |  |  | 2 |  |
| 19-Nov-23 | 18 |  |  |  |  |  |  |  |  |  |
| 10-Feb-24 | 12 | 4 | 3 | 4 |  |  |  |  |  |  |
| 6-May-24 | 18 | 2 | 6 | 3 |  |  |  |  |  |  |
| 26-Jun-24 | 45 | 4 | 7 | 6 | 4 | 4 | 2 | 1 | 1 | 2 |
| 7-Sep-24 | 21 | 6 | 1 | 3 |  |  |  |  |  |  |
| 7-Dec-24 | 29 |  | 3 | 4 | 2 |  |  |  |  |  |

### Survey of Sympatric Reservoirs at Affected and Unaffected Sites

To evaluate broader biogeographic patterns, we sampled additional coastal sites in Panama—Galeta (9.40°N, 79.86°W) and Taboga (8.78°N, 79.53°W)—and at Réunion Island, France (Port Sainte-Rose, 21.12°S, 55.79°E; Étang-Salé, 21.26°S, 55.33°E). Swabs were collected in August 2023 (Panama) and December 2023 (Réunion) following the protocol described in (7). In Panama, samples were stored at –80°C for approximately eight months prior to shipment at room temperature. Samples from Réunion were shipped in RNAlater and processed immediately upon arrival at Cornell University. In total, 44 specimens were collected from Galeta (Panama), 25 from Taboga (Panama), and 35 from Réunion (France) (Table S2). Swabbed substrates varied among sites depending on local benthic assemblages.

### DNA Extraction

All swab samples (from Grassy Key, Panama, and Réunion) were processed identically. Swabs were removed from RNAlater, transferred into Bead Basher tubes (Zymo Research), and homogenized by bead beating as part of the Quick-DNA Tissue/Insect Miniprep Kit (Zymo Research), followed by extraction per the manufacturer’s instructions.

For planktonic samples, thawed GF/D filters were sectioned with heat-sterilized razor blades, and half of each filter was extracted using the same Zymo protocol. DNA concentrations were quantified by the Quant-iT PicoGreen dsDNA Assay Kits (Invitrogen) using an ABI StepOne Real-Time PCR instrument.

### DaScPc Detection

DaScPc presence was determined using a combination of quantitative PCR (qPCR), nested PCR, and phylogenetic validation.

#### Quantitative PCR (qPCR)

Each reaction contained 1 µL of template DNA, 1× SSO Probes Supermix (Bio-Rad), and 200 nM each of primers Phil-28SF and Phil-28SR and probe Phil-28S-Pr (FAM-TAMRA labeled), targeting the DaScPc 28S rRNA gene (7, 8). Duplicate reactions per sample were compared against 28S rRNA oligonucleotide standards (10–10⁸ copies per reaction). Samples were considered positive when both replicates exceeded the detection threshold (>10² copies per reaction) based on the detectability of the oligonucleotide standards (Fig. 1) and (7).

**Figure 1:**
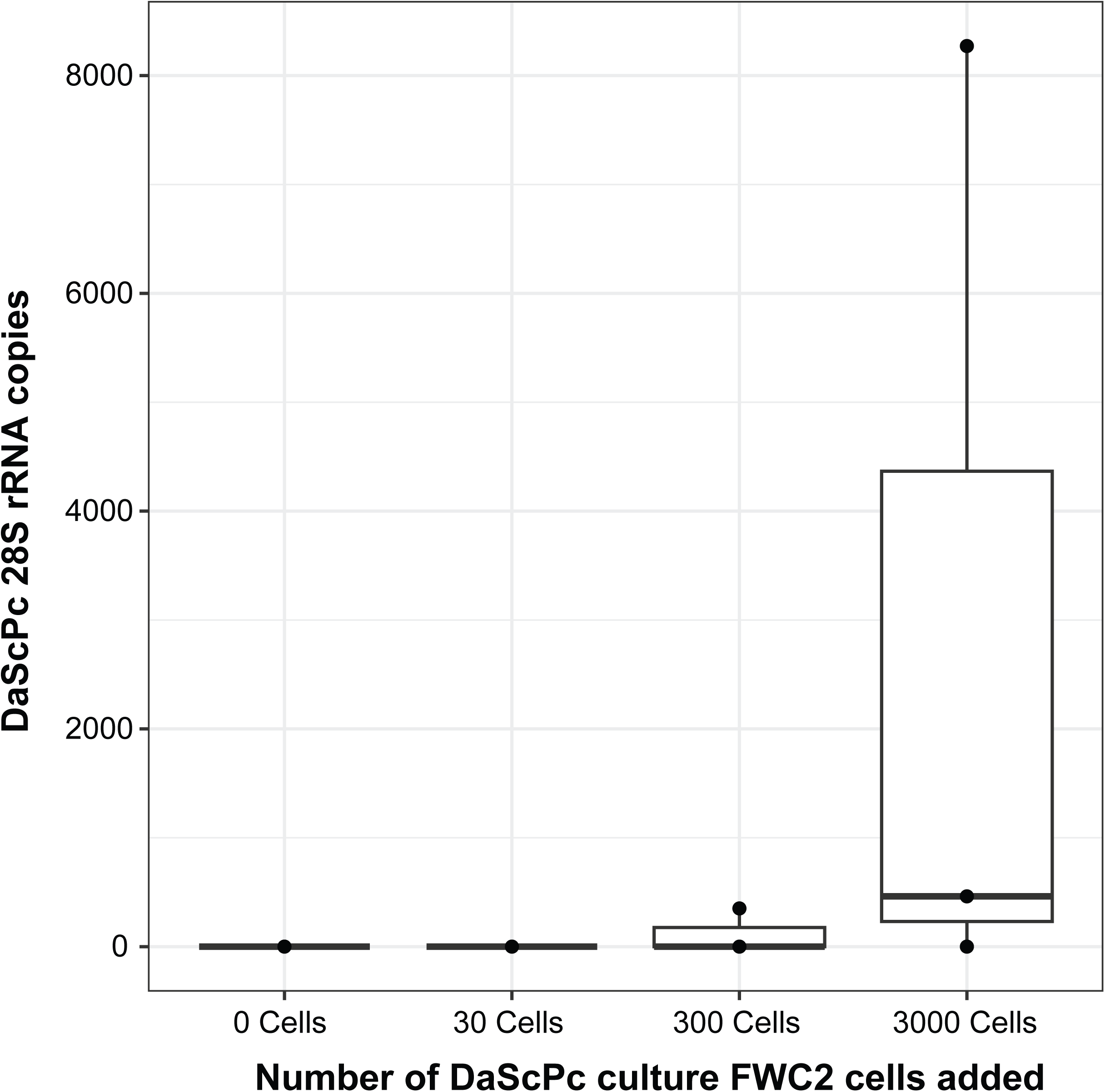
Assessment of swab surface recovery. Zero to 3000 DaScPc culture FWC2 cells were amended on Astrangia poculata in separate aquaria, then swabbed according to field protocols. The abundance of DaScPc 28S rRNA genes was determined by qPCR. Boxes illustrate the interquartile range (IQR), with whiskers indicating data extremes defined by 1.5 × IQR, and the median shown as a horizontal line within each box.

#### Nested PCR and Sanger Sequencing

All qPCR-positive samples were amplified using primers 384F/1147R (25), followed by a nested reaction with primers scutico-634F (7) and 1147R. Amplicons were visualized on agarose gels, purified with Clean & Concentrator-5 columns (Zymo Research), and sequenced at the Cornell Biotechnology Resource Center.

#### Phylogenetic Analyses

Sequences were trimmed and compared to the NCBI non-redundant (nr) database using BLASTn (26). Sequences with >92% identity to known scuticociliates were aligned with reference taxa using MUSCLE (Edgar 2004). Phylogenetic reconstruction employed neighbor-joining analysis with the Kimura 2-parameter model and gamma-distributed rate variation in MEGA X (27). Samples clustering within the DaScPc FWC2 clade were classified as DaScPc-positive; those clustering within the *Acropora*/CHN/2009 lineage (HM030718.1, HM030719.1) were designated as *Philasteridae* sp. (28).

### Estimation of Macroalgal Cover, algal identification, and Coral Proximity

GoPro photographs of swabbed sites were analyzed to estimate macroalgal cover and proximity to coral tissue. Ten images per sampling event were randomly selected and scored according to algal contact: (1) overlapping coral, (2) within 5 cm but not touching, or (3) >5 cm away or absent. Percent algal cover was calculated for all algae combined and by individual morphotypes where discernible. Dominant algae from December 2024 were collected for taxonomic identification by snorkeling. Briefly, algal pieces were clipped and placed into Whirl-Pak bags (Whirl-Pak) until processing. After surveying, algal pieces were placed into 5 mL cryovials (VWR), placed in 2 mL of RNAlater, and immediately frozen until further processing. Snippets were extracted using the Quick-DNA Fecal/Soil Miniprep Kit (Zymo Research) and amplified with eukaryotic 18S rRNA primers eukA and eukB (29). Amplicons were Sanger-sequenced and taxonomically classified via BLASTn searches against the NCBI nr database. Phylogenetic reconstruction was done first by aligning the 18S rRNA gene (MUSCLE (53)), then using neighbor joining and the Kimura-2 model of substitution and gamma-distributed sites in MEGAX (27).

### Statistical Analyses

All statistical analyses were performed in R (v4.3.1) (30). Data visualization employed the package ggplot2 (31, 32). Nonparametric comparisons between sample groups were conducted using Wilcoxon rank-sum tests implemented in the stats package (v3.6.2). Linear regression lines were generated using *geom_smooth* and the method *lm* within the *ggplot* function (31, 32).

## Results

### Swab Recovery Efficiency

Of the twelve *Siderastrea siderea* fragments incubated with the DaScPc culture FWC2, three produced positive qPCR amplicons (Fig. 1). The mean 28S rRNA copy number across the culture dilution series was 3.33 ± 0.37 × 10⁵ copies cell⁻¹ during exponential growth. In contrast, total copies detected from coral fragment swabs ranged between 0.10 and 2.48 × 10⁵, indicating that only single cells or cell fragments were recovered. Because ribosomal copy number can vary substantially between cultured and environmental cells (33), cell-based quantification was not attempted. Nonetheless, these data suggest that field-based swab surveys likely underestimated the true environmental abundance of DaScPc.

### Environmental Conditions at Grassy Key

Environmental monitoring at Grassy Key spanned from August 2023 to December 2024. Water temperature showed strong seasonality, ranging from winter lows near 21 °C to summer highs above 33°C. Salinity remained stable (33–36), consistent with euhaline conditions. pH was relatively constant (8.3–8.4) through mid-2024 but declined to 7.65–7.93 in late 2024. Dissolved oxygen (DO) remained supersaturated (117–167%), likely reflecting persistent wind-driven mixing (Fig. 2; Table S1).

**Figure 2:**
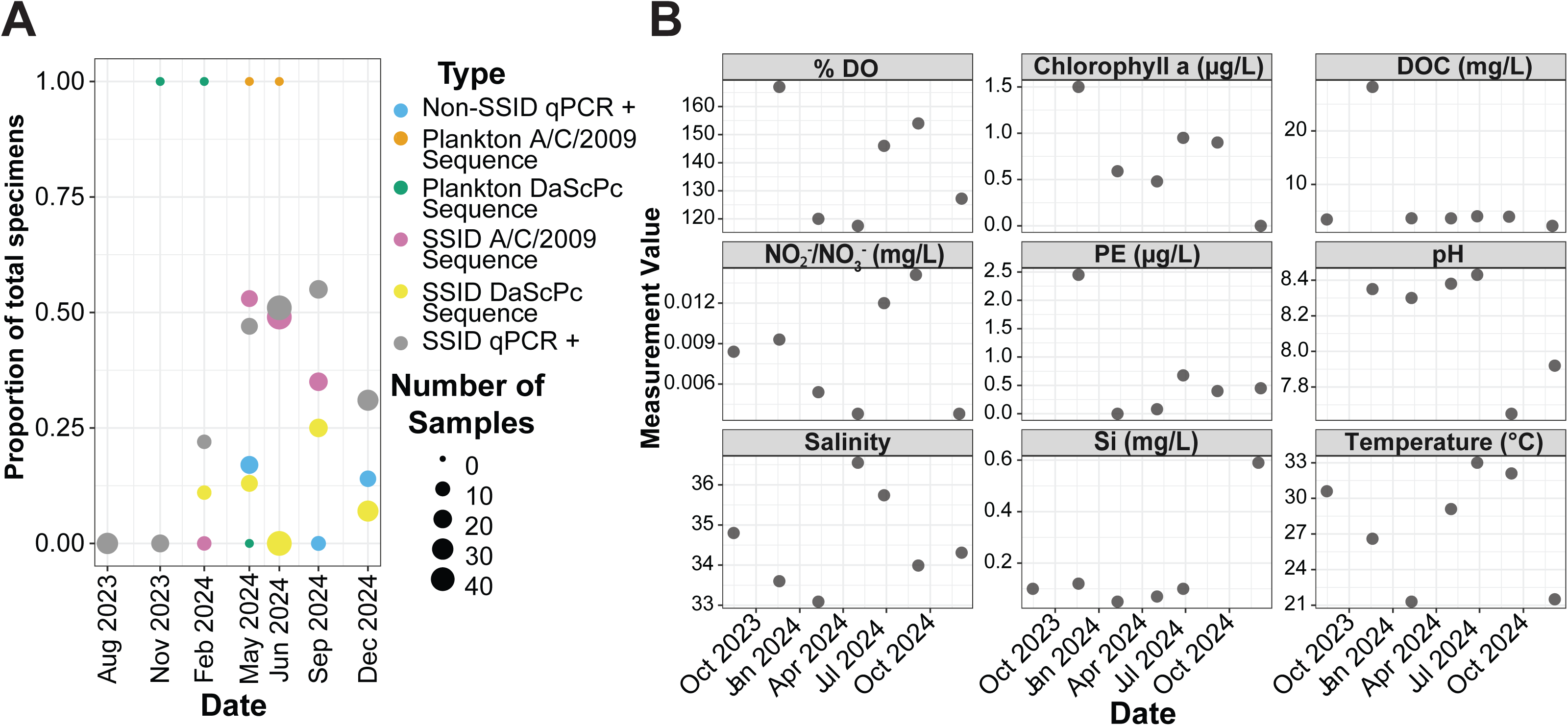
Temporal variation in scuticociliate prevalence and environmental characteristics. (A) Proportion of swabs that generated DaScPc 28S rRNA amplicons by qPCR in *Siderastrea sidereal (SSID)* and other surfaces. qPCR positive sequences were then subject to screening using nested conventional PCR; sequences that matched DaScPc and the related ciliate *Acropora*/CHN/2009 (A/C/2019) were expressed as a proportion of total swabs; (B) Physicochemical parameters and phytoplankton pigments (Chlorophyll a and phycoerythrin (PE)) were measured at the time of sampling.

Chlorophyll *a* followed a seasonal trend, with lowest concentrations in winter and peaks between spring and fall. The highest recorded value occurred in November 2023 (1.5 µg L⁻¹), coinciding with elevated phycoerythrin (2.55 µg L⁻¹; Fig. 2). A pronounced DOC spike (28 mg L⁻¹) was also observed that month, consistent with storm-related runoff and sediment resuspension documented in field notes (Table S1). Outside this event, DOC remained below 4 mg L⁻¹. Dissolved inorganic nutrients (NO₂⁻ + NO₃⁻ and Si) were generally low (0.003–0.012 mg L⁻¹), while NO₃⁻ and PO₄³⁻ were below detection limits throughout the study period (Table S1).

### Temporal Dynamics of DaScPc at Grassy Key

A total of 65 swab specimens collected between August 2023 and December 2024 yielded 28S rRNA qPCR amplicons, the vast majority of which (n = 60) were collected from *Siderastrea siderea*, while the rest were from turf algae (Table S2). No positive detections were obtained from sponge, seagrass, or other substrates.

Sequencing of qPCR-positive amplicons identified 10 specimens clustering with DaScPc, 44 with the *Acropora*/CHN/2009 clade, and one unrelated ciliate (Fig. 3). DaScPc was first detected in February 2024 and persisted intermittently through December 2024, absent only in June 2024. Plankton samples yielded qPCR amplicons in four months, but 18S rRNA sequencing confirmed DaScPc in only two instances (November 2023 and February 2024). An additional plankton amplicon from June 2024 clustered with *Acropora*/CHN/2009 (Fig. 2).

**Figure 3:**
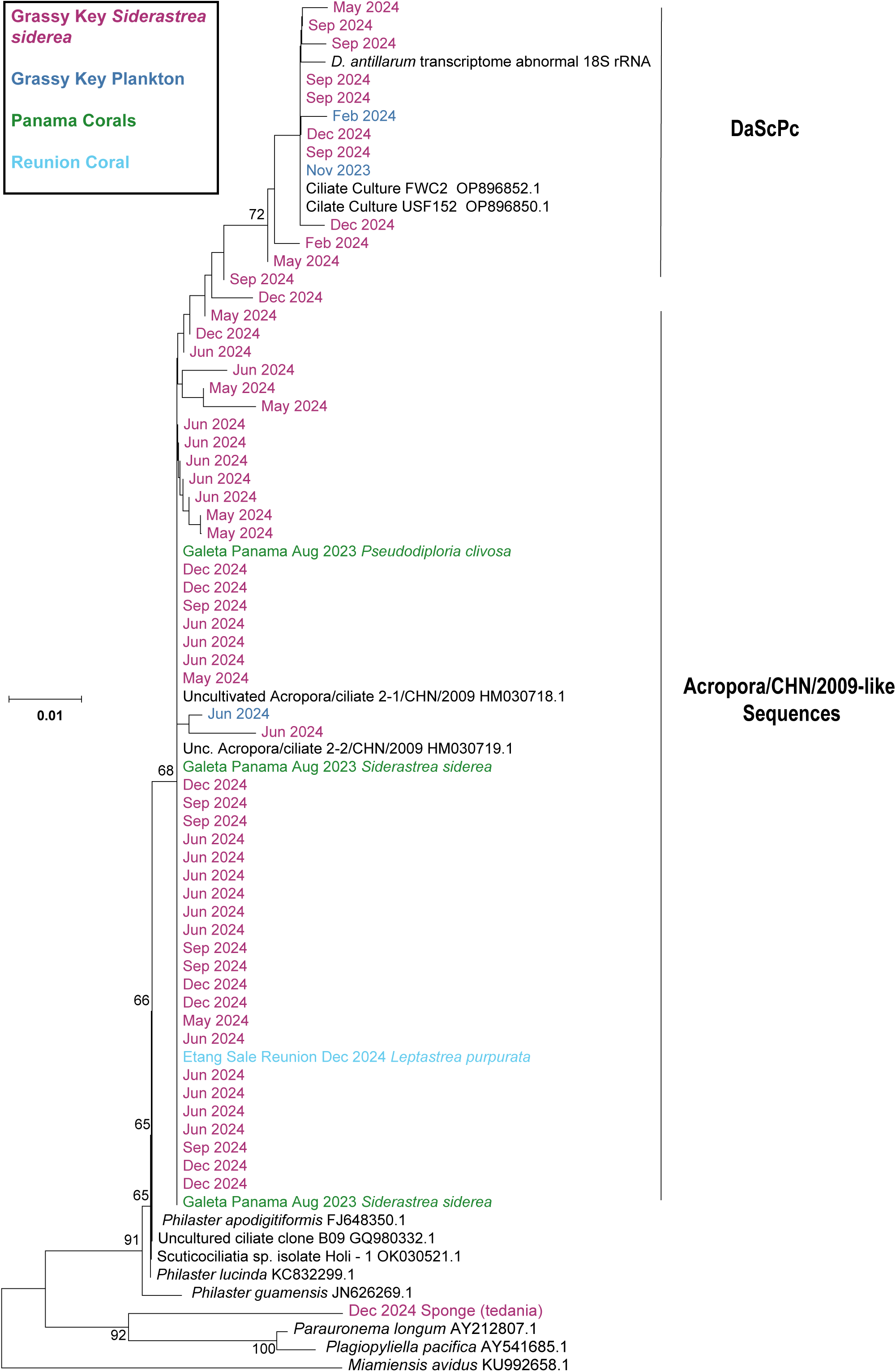
Phylogenetic reconstruction of scuticociliate sequences from swabs collected from Siderasterea siderea from Grassy Key (purple), coral from Panama (green), and coral from Reunion (teal). Scuticociliate sequences from planktonic samples collected at Grassy Key are blue. The reconstruction was performed based on a 299 nt alignment (MUSCLE (53)) by neighbor-joining, employing the Kimura-2 model and gamma substitution model for sites in MEGAX with 100 bootstraps (27).

Median qPCR copy numbers did not differ significantly among DaScPc-positive, *Acropora*/CHN/2009-positive, and other ciliate sequences (Wilcoxon P > 0.05; Fig. 4). Correlations between DaScPc prevalence and measured environmental parameters were not statistically significant. However, qualitative trends were observed: DOC and chlorophyll *a* both peaked in November 2023, coincident with the first DaScPc detection in the planktonic fraction, while *Acropora*/CHN/2009 prevalence increased through spring 2024 and declined after summer (Fig. 2).

**Figure 4:**
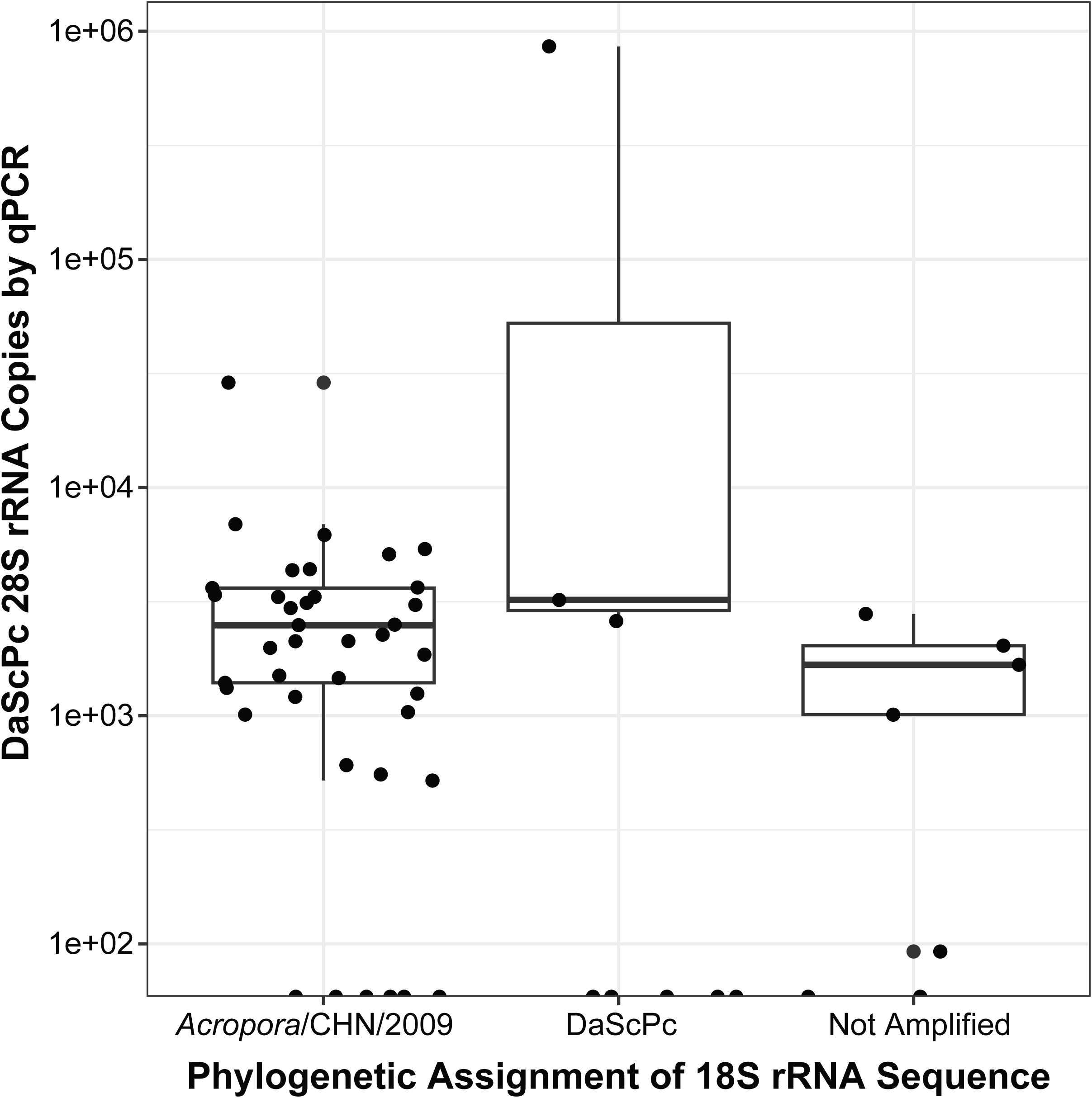
Comparison between DaScPc 28S rRNA quantity (determined by qPCR) and phylogenetic affiliation of swab-derived 18S rRNA sequences (by conventional PCR). qPCR quantities in swabs that did not amplify by 18S rRNA conventional PCR, yet yielded qPCR detections are presented for comparison. Boxes illustrate the interquartile range (IQR), with whiskers indicating data extremes defined by 1.5 × IQR, and the median shown as a horizontal line within each box.

### Macroalgal Associations

Dominant macroalgae at the site were identified by 18S rRNA sequencing as *Laurencia filiformis*, *Spyridia filamentosa*, *Dasya hutchinsiae*, *Asparagopsis taxiformis*, and the green alga *Caulerpa venticellata* (Fig. 5). Algal proximity scores did not differ significantly between DaScPc-positive and negative corals (P > 0.05; Fig. 6). Nevertheless, seasonal fluctuations in percent cover of *L. filiformis* corresponded with temporal variation in both DaScPc and *Acropora*/CHN/2009 detection (Fig. 7), suggesting potential indirect links between algal bloom dynamics and ciliate occurrence (Fig 7B; *Acropora*/CHN/2009 r=0.6, DaScPc r=0.63).

**Figure 5:**
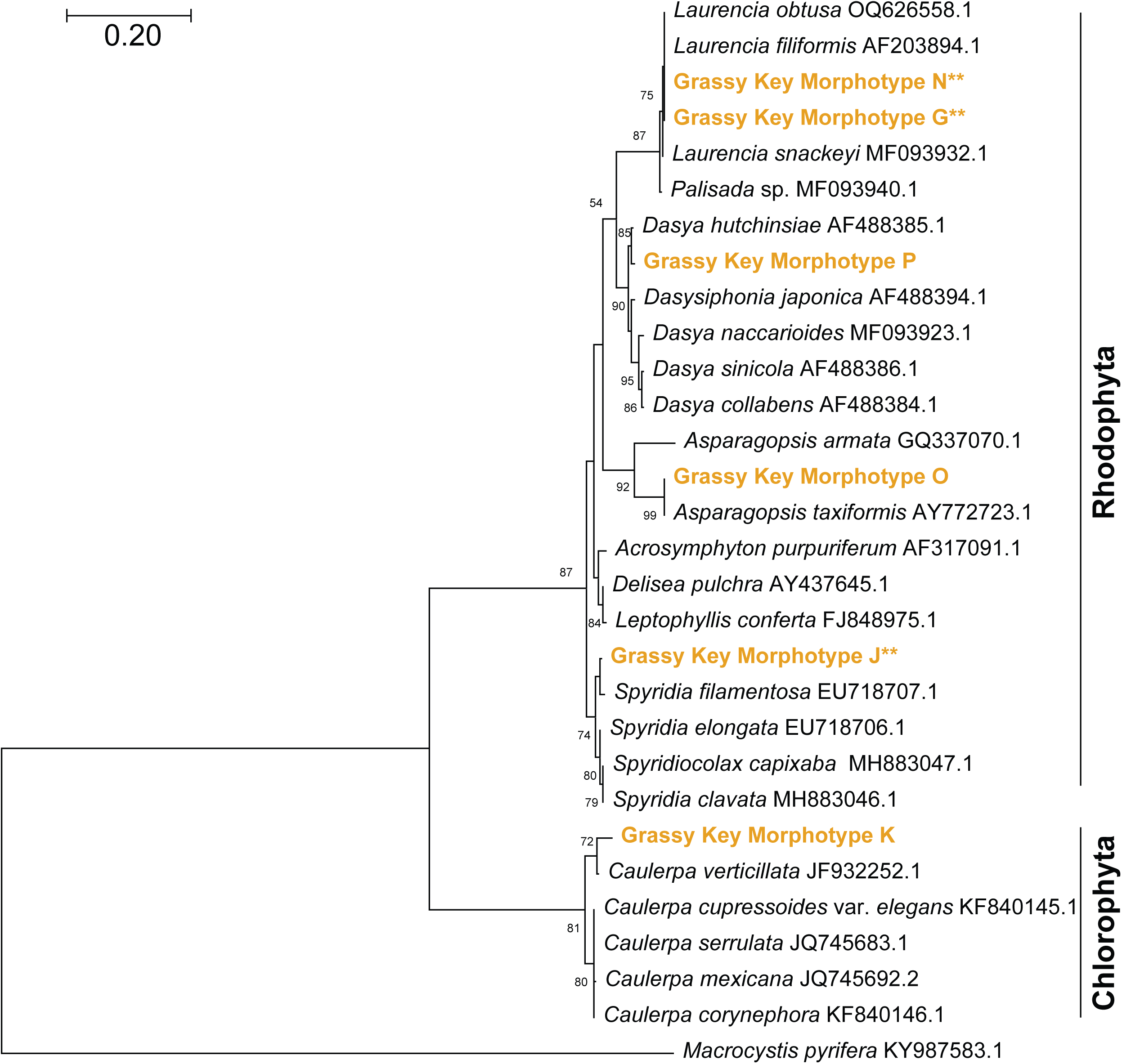
Phylogenetic representation of dominant macroalgae from Grassy Key (orange). The reconstruction is based on a 680 nt alignment of the 18S rRNA gene (MUSCLE (53)) using neighbor joining and the Kimura-2 model of substitution and gamma distributed sites in MEGAX with 100 bootstraps (27). Samples collected from macroalgae during the observed May and December 2024 blooms are designated with two astrixs (**).

**Figure 6:**
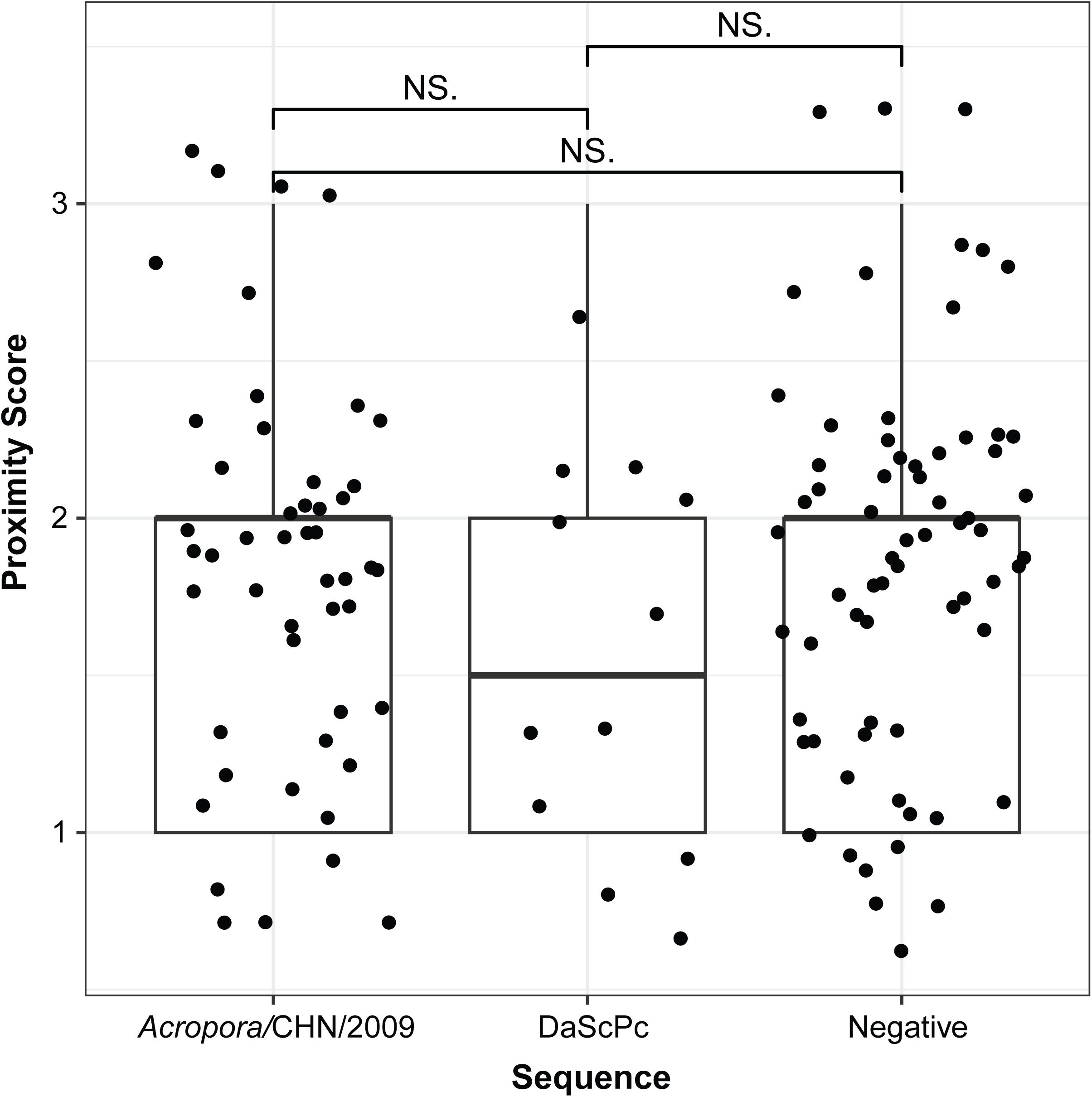
Detection of DaScPc and Acropora/CHN/2009 in relation to macroalgal proximity. Proximity was scored from underwater photographs (1) if the algae were touching or overlapping swabbed corals; (2) were not touching but within ∼5cm of corals; (3) were more than 5cm from corals. Significance was determined using a Wilcoxon signed-rank test. Boxes illustrate the interquartile range (IQR), with whiskers indicating data extremes defined by 1.5 × IQR, and the median shown as a horizontal line within each box.

**Figure 7:**
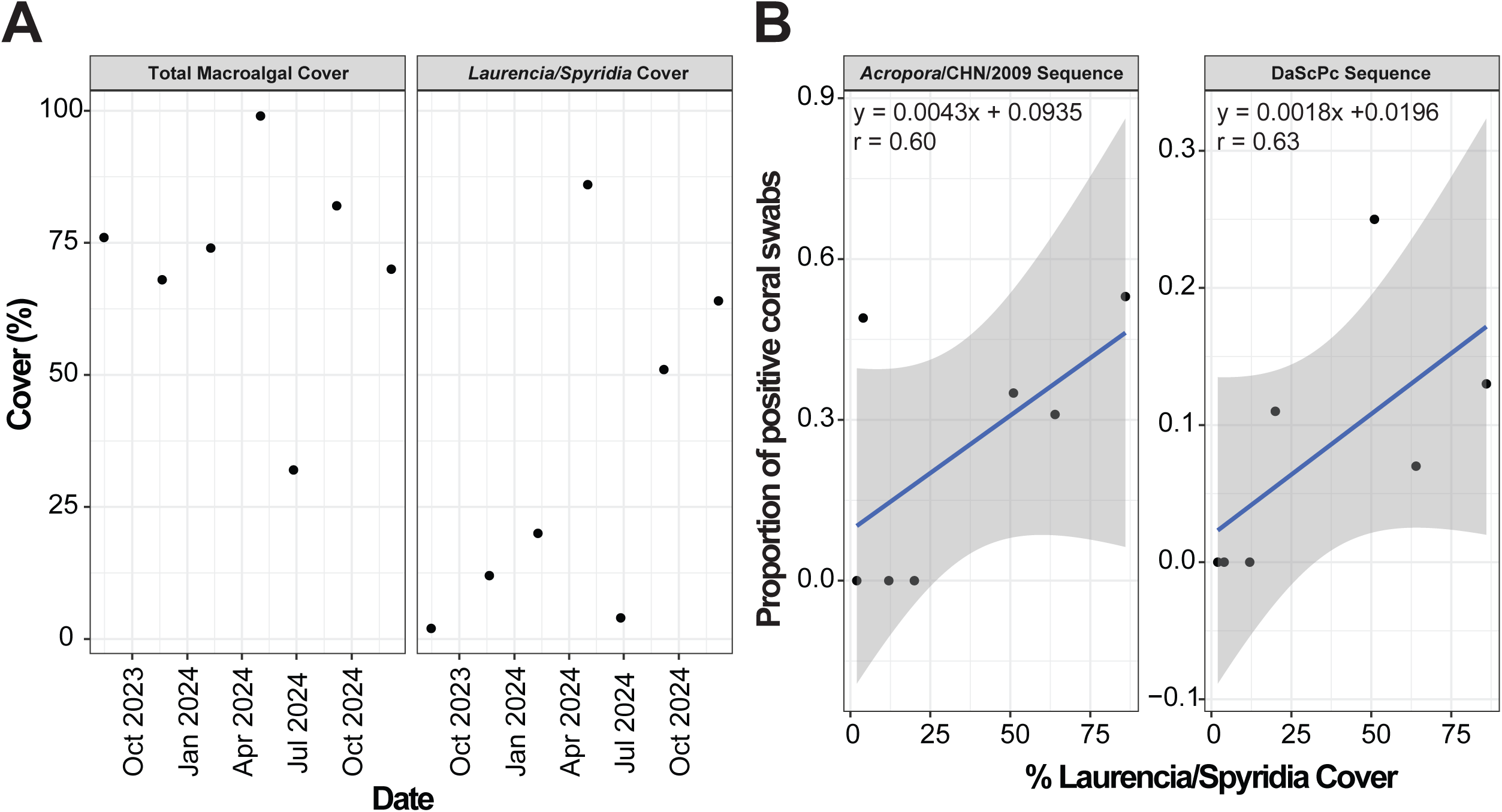
(A) Percent macroalgal cover and percent *Laurencia/Spyridia* cover determined from underwater photographs over time. Cover was estimated from photographs (n = 10 per sampling time) by randomization and manual scoring (B) Regression of (left) DaScPc 18S rRNA sequence positive proportion and (right) *Acropora*/CHN/2009 18S rRNA sequence positive coral swabs with % Laurencia/Spyridia cover. The gray zone is the 95% confidence interval.

### Surveys from Réunion and Panama

Swab surveys from Réunion and Panama yielded four qPCR-positive specimens: two *Siderastrea siderea* and one *Pseudodiploria labyrinthiformis* from Galeta (Panama), and one *Lepastrea purpurea* exhibiting black band disease from Étang-Salé (Réunion) (Table S1). Sequencing clustered all four sequences with *Acropora*/CHN/2009 rather than DaScPc, indicating the absence of confirmed DaScPc detections at either location (Fig. 3).

## Discussion

The finding that DaScPc, a ciliate responsible for the mass mortality of the keystone herbivore *Diadema antillarum*, can persist in marine environments independently of its known host is an important step in understanding its disease ecology. Ciliates are fundamental components of the microbial loop, serving as essential energy conduits in marine food webs (2, 3). However, this ecologically vital group also includes significant pathogens, whose environmental presence dictates the reemergence of marine diseases (34). Our positive detections confirmed by Sanger sequencing at Grassy Key (FL, USA), a site absent of *D. antillarum*, demonstrates that *DaScPc* is a natural presence in coastal and coral reef ecosystems, challenging the assumption that it is solely dependent on its urchin host. By utilizing molecular methods like qPCR and nested PCR, we were able to survey multiple putative environmental niches and improve detection confidence. The detection of *DaScPc* on coral surfaces, macroalgae, and in the water column shows that DaScPc can utilize multiple environmental reservoirs (7). This finding supports the emerging consensus that the environmental persistence and dispersal of protistan pathogens are critical, often cryptic, factors in disease dynamics (34, 35), and establishes a foundation and key considerations for effective disease monitoring and reef conservation.

Our molecular surveillance provides crucial insights into the ecological niche of DaScPc, highlighting a strong benthic association and potential dependence on primary production. The increased detection of DaScPc on coral surfaces and in the plankton fraction, often co-occurring with the closely related ciliate *Acropora*/CHN/2009 (28), suggests a presence in the benthic microbial community that does not necessarily equate to host disease. This observation aligns with the dual role of ciliates in marine systems, both as trophic links and potential cryptic pathogens (3, 34).

The temporal patterns we observed, where DaScPc prevalence was associated with periods of benthic primary production such as macroalgal blooms, supports the Dissolved Organic Carbon (DOC), Disease, Algae, and Microbes (DDAM) model (21). The DDAM model posits that DOC released by fleshy macroalgae drives the proliferation of opportunistic microbes, including potential pathogens. Our finding that DaScPc was observed during these productive periods, even without association with host disease, suggests the ciliate may be an opportunistic heterotroph that leverages these nutrient pulses (36, 37). Detection in the plankton fraction during the less productive fall and winter seasons is likely to reflect resuspension from these benthic surfaces (38) or transient growth on free-living microbial prey, rather than suggesting that sediments are a primary reservoir.

Our geographic findings emphasize the complexities of marine pathogen epidemiology, particularly the concept of the environmental reservoir. The positive detections at Grassy Key confirm the ability of DaScPc to persist in the absence of the host (D. *antillarum*), underscoring the potential role of benthic substrates in disease maintenance. This is supported by the high detection rates on *Siderastrea siderea*, corroborating earlier work that suggested corals can act as reservoirs for DaScPc (Hewson et al., 2023; Vilanova-Cuevas et al., 2025b). However, the absence of DaScPc from sites in Panama and Réunion, where previous outbreaks were suspected, highlights spatial patchiness in its distribution or an opportunistic lifestyle.

This sporadic detection aligns with the predicted dynamics of protistan pathogens that may follow a “boom and bust” lifecycle (39–43). Under this model, the pathogen maintains a low, cryptic environmental abundance (a ’bust’ phase) that is difficult to detect, interspersed with rapid population explosions (a ’boom’ phase) often triggered by environmental shifts. The lack of detection outside our focal site may be due to this low cell abundance, suggesting a conservation of detection effort is warranted when surveying the environmental pool of a potentially rare or transient protist (40). The unresolved question of whether DaScPc is a stable component of the coral microbiome or primarily a transient environmental occupant remains an area for future research.

The lack of a clear, instantaneous relationship between DaScPc prevalence and measured physicochemical factors (e.g., salinity, instantaneous temperature) suggests that the ciliate’s occurrence may be driven less by short-term water conditions and more by broader, chronic environmental disturbances or productivity shifts (21, 40, 43). This aligns with the complex ecological landscape of microbial pathogens, where multiple stressors often interact to facilitate disease outbreaks (44–46). Thermal tolerance is a major constraint for many marine organisms (47–50), and extreme events like the 2023 Caribbean heatwave could limit DaScPc abundance (51, 52). However, our positive detections in subsequent high-temperature periods indicate that temperature alone may not be the sole determinant of its presence. Instead, factors such as nutrient loading, shifts in the microbial food web, and host density likely interact to dictate whether the ciliate maintains a stable reservoir population or initiates an outbreak (8). Future work should focus on longitudinal, high-frequency sampling to better link these broad environmental shifts with changes in DaScPc’s environmental abundance and subsequent disease emergence.

### Limitations

Several methodological and logistical limitations must be considered when interpreting these results. Swab-based qPCR detection requires a minimum cell threshold, meaning non-detections cannot be taken as evidence of true absence. Confirmed positives via Sanger sequencing provide stronger support for presence but are limited by sample size. Sample preservation, nucleic acid extraction efficiency, and potential cell loss in RNALater may have influenced detection rates. Additionally, qPCR can produce false positives, emphasizing the need for sequencing-based confirmation in environmental surveys.

Spatial and temporal coverage was limited, potentially underestimating the full ecological distribution of *DaScPc*. Limited detections at sites with previous putative outbreaks suggest that the ciliate may be environmentally passive in some contexts, underscoring the need for expanded, longitudinal surveys. While corals were the primary positive substrate, it remains unclear whether *DaScPc* is a stable component of the coral microbiome or functions primarily as a transient occupant within reef habitats.

## Conclusions

This study shifts the understanding of DaScPc by demonstrating that this ciliate is a naturally occurring component of the coral reef environment, capable of environmental persistence independent of its host, *Diadema antillarum* (7). Molecular detection on benthic surfaces and in the plankton fraction, particularly at urchin-free sites, confirms that corals and macroalgae serve as crucial natural reservoirs, supporting the broader epidemiological concept that protistan pathogens maintain populations between host-mortality events. Our findings reveal a critical link between DaScPc’s presence and the microbial food web. Detection during macroalgal blooms is consistent with the DDAM model, suggesting that environmental productivity and dissolved organic carbon availability may be key factors governing population dynamics. This environmental association, along with the co-occurrence of the non-pathogenic ciliate *Acropora*/CHN/2009, underscores the ecological complexity of this microbial group and the need to distinguish a ciliate’s general ecological role from its pathogenic potential.

The demonstrated ecological flexibility of DaScPc implies that its parasitic interactions are intermittent and governed by poorly constrained environmental or host-related circumstances. This necessitates a transition to high-resolution surveillance. Future research must focus on clarifying the feeding ecology of DaScPc and the specific environmental triggers that govern its shift from an innocuous reservoir state to an outbreak phase. Expanded, longitudinal surveys of macroalgal blooms, nutrient fluxes, and temperature anomalies are essential to forecasting ciliate distribution and mitigating disease reemergence in vulnerable coral reef ecosystems.

## Acknowledgements

A portion of this work was funded by the Simons Foundation Marine Microbiology Postdoctoral Fellowship and the Department of Biological Sciences and College of Liberal Arts and Sciences at Northern Illinois University to M.W.H. This work was supported by NSF award OCE-2049225 awarded to I.H., and NSF award OCE- awarded to I.H. and M.W.H. Sampling was performed at Grassy Key under permit SAL-23-2564-SR from the Florida Fish and Wildlife Conservation Commission. Samples from Panama were collected under permit ARBG-073-2022 from the Ministerio de Ambiente, Panama. Samples from Réunion were collected with permission from the Service Eau & Biodiversité, Direction de l’environnement de l’aménagement et du logement, Prefet de la Région Réunion.

## Data Availability

Physicochemical characteristics at the Grassy Key site are provided in Supplemental Table 1. qPCR detection and Sanger sequence identities after provided in Supplemental Table 2. Sequence data have been deposited at NCBI GenBank under accessions PV505242-PV505300.

